# Region-Specific Delta and Alpha2 Relative Power Alterations in Male Adolescent Schizophrenia: Occipital Predominance in Resting-State EEG

**DOI:** 10.64898/2026.07.14.738386

**Authors:** Suvojit Hazra, Nilkanta Chakrabarti

## Abstract

Resting-state electroencephalography (EEG) studies consistently report increased slow-wave activity and reduced alpha power in schizophrenia. However, the regional distribution of EEG relative power (RP) across finer frequency bands, particularly in adolescent schizophrenia, remains poorly characterized. We analyzed a publicly available resting-state EEG dataset comprising 45 male adolescents with schizophrenia (SCZ; mean age: 12.3 ± 1.2 years) and 39 age- and sex-matched healthy controls (CON; mean age: 12.3 ± 1.3years), recorded using 16 scalp electrodes (10–20 system). Following band-pass filtering, artefact subspace reconstruction, spline interpolation, average re-referencing and independent component analysis, RP was computed for nine frequency bands across five cortical regions (frontal, central, parietal, temporal, and occipital). Group differences were assessed using three-way repeated-measures analysis of variance (ANOVA), followed by Bonferroni false discovery rate (FDR)-corrected post-hoc pairwise comparisons using estimated marginal means (emmeans). Compared to controls, the SCZ group showed significantly higher delta RP in the frontal, parietal, and occipital regions and significantly lower alpha2 RP, dissociable from alpha1, in the central, parietal, temporal, and occipital regions (all FDR < 0.001). The occipital cortex showed the greatest increase in delta RP (|Cohen’s *d*|=0.82) and the greatest reduction in alpha2 RP (|*d*|=0.73), with moderate-to-large effect sizes. Topographic maps demonstrated widespread delta enhancement and attenuated occipito-parietal alpha2 activity in the SCZ group. These findings demonstrate region-specific alterations in resting-state EEG relative power in adolescent schizophrenia, particularly within the occipital cortex, and suggest that regional RP may serve as a promising neurophysiological feature for early-onset schizophrenia.

**Highlights:**

- Resting-state EEG relative power was analyzed in adolescent schizophrenia.
- Delta relative power increased (occipital > parietal > frontal).
- Alpha2, but not alpha1, relative power decreased (occipital > parietal > temporal > central).
- Occipital cortex showed the largest EEG spectral abnormalities.
- Nine-band analysis improved regional spectral characterization.

## 1. Introduction

Schizophrenia is a chronic, debilitating neuropsychiatric disorder affecting approximately 1% of the global population, characterized by positive symptoms (hallucinations, delusions), negative symptoms (affective flattening, avolition), and cognitive deficits. Although onset typically occurs in early adulthood, prodromal features are frequently detectable in adolescence, making early neurobiological characterization a priority for timely intervention^1^. During the prodromal stage, working-memory impairment is associated with aberrant activation of the dorsolateral and ventrolateral prefrontal cortices and anterior cingulate cortex, together with reduced frontoparietal connectivity, indicating early disruption of executive neural circuits in adolescent schizophrenia^2^.

The electroencephalogram (EEG) provides a direct, millisecond-resolution window into cortical oscillatory dynamics and has been extensively studied in schizophrenia. A meta-analysis of spectral EEG abnormalities documented consistent patterns of increased delta and theta power alongside decreased alpha power in patients, with frontal and temporal scalp regions most frequently implicated^3^. These spectral aberrations have been interpreted within frameworks of thalamocortical dysrhythmia, impaired cortical inhibition mediated by parvalbumin-positive interneurons, and dysregulation of the excitation/inhibition (E/I) balance^4,5^.

Although resting-state EEG abnormalities in schizophrenia have been extensively investigated, several important gaps remain. Most previous studies analyzed absolute power or broad frequency bands (delta, theta, alpha, and beta), making it difficult to determine whether abnormalities are confined to specific sub-bands. In addition, previous relative power studies have reported heterogeneous findings because of differences in frequency definitions, analytical approaches, recording conditions, and cortical regions examined. Conventional four-band analyses described reduced occipital alpha activity in adult schizophrenia and schizoaffective disorder^6,7^, whereas later studies reported broader regional abnormalities involving frontal, central, parietal, and temporal cortices^8,9^. However, few studies have systematically evaluated relative power across multiple frequency sub-bands and cortical regions in adolescent schizophrenia using a rigorous statistical framework. Consequently, the regional distribution and frequency specificity of resting-state EEG abnormalities during early-onset schizophrenia remain incompletely characterized.

The present study addresses these gaps by characterizing resting-state EEG RP across nine frequency bands and five cortical regions in a well-characterized male adolescent schizophrenia cohort using a publicly available dataset^10,11^. Relative power was selected because it normalizes inter-individual differences in overall EEG amplitude and provides a measure of spectral redistribution that is less influenced by non-neural factors than absolute power^12,13^. We applied a comprehensive three-way repeated-measures ANOVA followed by multiple comparison correction to identify frequency- and region-specific alterations and generated topographic maps to visualize their spatial distribution. By combining high-resolution spectral decomposition with quantitative regional analysis, this study aimed to provide a more comprehensive characterization of resting-state oscillatory abnormalities in adolescent schizophrenia than has been available from previous broad-band EEG investigations.

## 2. Materials and methods

### 2.1 Participants and EEG Dataset

We utilized the publicly available male adolescent schizophrenia EEG dataset from Moscow University (http://brain.bio.msu.ru/eeg_schizophrenia.htm), comprising 45 individuals with a ICD-10 diagnosis of schizophrenia (SCZ; mean age 12.3 ± 1.2 years) and 39 matched healthy controls (CON; mean age 12.3 ± 1.3 years) ^10,11^. All participants were recruited and assessed by clinicians at the National Center of Mental Health of the Russian Academy of Medical Sciences. Diagnostic criteria, inclusion and exclusion criteria, and experimental assay protocols are described in detail in the original dataset publication^10,11,14^. All patients were not on medication at the time of recording^10,11^.

### 2.2 EEG Recording

Resting-state, eyes-closed EEG was recorded from 16 scalp electrodes arranged according to the international 10-20 system, referenced to coupled ear electrodes: Frontal (F7, F3, F4, F8), Central (C3, Cz, C4), Parietal (P7, P3, Pz, P4, P8), Temporal (T7, T8), and Occipital (O1, O2). Electrode labels were updated from the legacy convention (T3, T4, T5, T6) to the Modified Combinatorial Nomenclature (MCN) system ^15^ (**Figure 2j**). The sampling rate was 128 Hz and recording duration was 60 seconds per participant (7,680 samples total)^10,11^.

### 2.3 EEG Pre-processing

Raw EEG data (.eea format) were imported into EEGLAB (v2021.0)^16^ running under MATLAB 2019a. A Hamming-windowed finite impulse response (FIR) bandpass filter (0.5-50 Hz; transition bandwidth 0.5 Hz; cutoff range 0.25-50.25 Hz) was applied. Artefact subspace reconstruction (ASR) was performed using the clean_rawdata() plugin^17,18^ with the following parameters: flat-line removal threshold >10 seconds; minimum channel correlation ≥0.70; ASR threshold = 20 with a 0.5-second rejection window; maximum allowable power ratio = 3.5; maximum out-of-bound channel threshold = 25%. Following artefact removal, rejected channels were interpolated by spline interpolation and data were re-referenced to the average. Extended infomax independent component analysis (ICA) was applied via runica^19^, and brain-relevant independent components were identified and cleaned using the ICLabel plugin^20^ combined with a Wavelet-ICA approach to zero out artifactual temporal segments (z-score ≥ ±2 within 1-second windows).

### 2.4 Frequency Bands

Nine frequency bands were defined according to the schizophrenia EEG literature^3^: (1) delta (0.5-3.5 Hz), (2) theta1/low-theta (4-5.5 Hz), (3) theta2/high-theta (6-7.5 Hz), (4) alpha1/low-alpha (8-10 Hz), (5) alpha2/high-alpha (10.5-12.5 Hz), (6) beta1/low-beta (13-16 Hz), (7) beta2/moderate-beta (17-23 Hz), (8) beta3/high-beta (24-30 Hz), and (9) gamma (31-50 Hz).

### 2.5 Relative Power Computation

Mean absolute power spectral density (AP) was computed for each frequency band using the Fast Fourier Transform (FFT) via EEGLAB’s spectopo() function, employing 2-second Hamming-windowed segments (256-point FFT; resolution 0.5 Hz). A natural log transform was applied to linearize the power values. Individual electrode AP values were averaged within each of five cortical regions (frontal, central, parietal, temporal, occipital). Relative power (RP) was then derived for each band as:

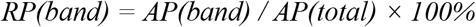

where AP(total) is the sum of AP across all nine bands.

### 2.6 Statistical Analysis

All statistical analyses were performed in R (v4.1.1)^21^. Normality and homogeneity of variance were assessed using Shapiro–Wilk and Levene tests, respectively, before proceeding with ANOVA. A three-way repeated-measures ANOVA (Analysis of Variance) was conducted for RP using anova_test() from the rstatix R package^22^, with Diagnosis (SCZ, CON) as a between-subject factor and Band (nine levels) and Region (five levels) as within-subject factors. A three-way interaction threshold of p < 0.001 was applied for significance. Significant ‘Diagnosis *x* Band’ interactions were followed by two-way repeated-measures omnibus ANOVAs for each band separately. Post-hoc pairwise comparisons between SCZ and CON across Band and Region were performed using the emmeans R package^23^ with Bonferroni false discovery rate (FDR) correction; p < 0.001 was the significance threshold. To quantify the magnitude of the differences, Cohen’s *d* effect sizes for the pairwise comparisons were calculated using the pooled standard deviation.

## 3. Results

### 3.1 Three-Way ANOVA for Relative Power

The three-way repeated-measures ANOVA for RP (**Table 1**) demonstrated a highly significant ‘Diagnosis *x* Band’ interaction (F(8,3690) = 23.147, p < 0.001, ηg^2^ = 0.048) and ‘Region *x* Band’ interaction (F(32,3690) = 24.864, p < 0.001, ηg^2^ = 0.177), indicating that the spectral composition of EEG activity and its regional distribution differed substantially between SCZ and CON across frequency bands. In contrast, the main effects of Diagnosis alone (F(1,3690) < 0.001, p = 0.998) and Region alone (F(4,3690) < 0.001, p = 1.000) were not significant, nor was the three-way ‘Diagnosis *x* Region *x* Band’ interaction (F(32,3690) = 0.554, p = 0.98). These findings indicated that diagnostic differences in RP were expressed as band- and region-specific spectral redistribution rather than as a uniform global shift.

**Table 1.** Summary of three-way repeated-measures ANOVA for EEG relative power (RP). Significance thresholds: * p < 0.001, ns = not significant.

| Effect | Dfn, Dfd | F-score | p-value | Significance | $\eta^2$ |
| --- | --- | --- | --- | --- | --- |
| Diagnosis | 1, 3690 | 0.00000387 | 0.998 | ns | 1.05e-09 |
| Region | 4, 3690 | 0.00000342 | 1 | ns | 3.71e-09 |
| Band | 8, 3690 | 1024.145 | <0.001 | * | 0.689 |
| Diagnosis × Region | 4, 3690 | 0.000004 | 1 | ns | 4.34e-09 |
| Diagnosis × Band | 8, 3690 | 23.147 | 6.89e-35 | * | 0.048 |
| Region × Band | 32, 3690 | 24.864 | 1.58e-131 | * | 0.177 |
| Diagnosis × Region × Band | 32, 3690 | 0.554 | 0.98 | ns | 0.005 |

### 3.2 Band-Wise Repeated-Measures ANOVA

Separate two-way repeated-measures omnibus ANOVAs were performed for each of the nine frequency bands (**Table 2**). Diagnosis exerted a significant main effect on RP in seven of nine bands (delta, theta1, theta2, alpha1, alpha2, beta2, and beta3; all p < 0.001), whereas no significant diagnostic effect was observed for the beta1 (p = 0.002) or gamma (p = 0.892) bands at the predefined significance threshold. Region exerted a significant main effect on RP in eight of the nine frequency bands (delta, theta1, theta2, alpha1, alpha2, beta1, beta2, and beta3; all p < 0.001), whereas no significant regional effect was observed for the gamma band (p = 0.945). Among the significant bands, the largest F-statistics for Diagnosis were observed in the alpha2 (F(1,3690) = 7827.963), delta (F(1,3690) = 5598.679), and alpha1 (F(1,3690) = 2579.013) bands indicating the strongest group differences in these spectral domains.

**Table 2.** Summary of band-wise two-way repeated-measures omnibus ANOVA for RP. Significance: * p < 0.001; ns = not significant.

| Band | Effect | Dfn, Dfd | F-score | p-value | Significance | $\eta^2$ |
| --- | --- | --- | --- | --- | --- | --- |
| Delta | *Diagnosis* | 1, 3690 | 5598.679 | <0.001 | * | 0.603 |
|  | *Region* | 4, 3690 | 5876.707 | <0.001 | * | 0.864 |
| Theta1 | *Diagnosis* | 1, 3690 | 1264.368 | 2.11e-238 | * | 0.255 |
|  | *Region* | 4, 3690 | 2028.892 | <0.001 | * | 0.687 |
| Theta2 | *Diagnosis* | 1, 3690 | 312.95 | 2.77e-67 | * | 0.078 |
|  | *Region* | 4, 3690 | 253.074 | 2.25e-192 | * | 0.215 |
| Alpha1 | *Diagnosis* | 1, 3690 | 2579.013 | <0.001 | * | 0.411 |
|  | *Region* | 4, 3690 | 9524.28 | <0.001 | * | 0.912 |
| Alpha2 | *Diagnosis* | 1, 3690 | 7827.963 | <0.001 | * | 0.680 |
|  | *Region* | 4, 3690 | 1211.842 | <0.001 | * | 0.568 |
| Beta1 | *Diagnosis* | 1, 3690 | 9.172 | 0.002 | ns | 0.002 |
|  | *Region* | 4, 3690 | 11.162 | 5.32e-09 | * | 0.012 |
| Beta2 | *Diagnosis* | 1, 3690 | 12.25 | 0.000471 | * | 0.003 |
|  | *Region* | 4, 3690 | 22.497 | 2.22e-18 | * | 0.024 |
| Beta3 | *Diagnosis* | 1, 3690 | 22.438 | 2.25e-06 | * | 0.006 |
|  | *Region* | 4, 3690 | 5.455 | 0.000223 | * | 0.006 |
| Gamma | *Diagnosis* | 1, 3690 | 0.018 | 0.892 | ns | 5e-06 |
|  | *Region* | 4, 3690 | 0.188 | 0.945 | ns | 0.000204 |

### 3.3 Post-Hoc Analysis: Significant Group Differences in RP

Post-hoc pairwise comparisons using estimated marginal means (emmeans) with Bonferroni false discovery rate (FDR) correction identified significant (FDR < 0.001) alterations in RP between the SCZ and CON groups in the delta and alpha2 frequency bands (**Figure 1, Table 3**).

**Table 3.** Post-hoc pairwise comparison of RP differences between CON (n=39) and SCZ (n=45) groups. Values are mean (SD, SEM) in %. Only significant band–region combinations reaching FDR < 0.001 are shown in Bold. Difference = SCZ mean ™ CON mean.

| Frequency Band | Region | Mean Relative power in % (SD, SEM) |  | Difference | FDR-adjusted p-value | \|Cohen's d\| |
| --- | --- | --- | --- | --- | --- | --- |
|  |  | CON (n = 39) | SCZ (n = 45) |  |  |  |
| **Delta** |  |  |  |  |  |  |
|  | **Frontal** | **28.83 (7.67, 1.23)** | **33.67 (11.30, 1.68)** | **4.84** | **0.00078** | **0.49** |
|  | Central | 22.06 (7.82, 1.25) | 25.34 (9.40, 1.40) | 3.28 | 0.022 | 0.38 |
|  | **Parietal** | **18.59 (6.72, 1.08)** | **23.87 (9.77, 1.46)** | **5.28** | **0.0002** | **0.62** |
|  | Temporal | 24.61 (7.36, 1.18) | 29.08 (12.15, 1.81) | 4.47 | 0.00191 | 0.44 |
|  | **Occipital** | **12.82 (5.43, 0.87)** | **19.65 (10.14, 1.51)** | **6.83** | **2.17e-6** | **0.82** |
| **Theta1** |  |  |  |  |  |  |
|  | Frontal | 15.32 (3.77, 0.60) | 17.16 (5.60, 0.83) | 1.84 | 0.201 | 0.38 |
|  | Central | 16.61 (5.21, 0.83) | 20.04 (6.33, 0.94) | 3.43 | 0.017 | 0.59 |
|  | Parietal | 13.91 (4.65, 0.74) | 15.93 (5.46, 0.81) | 2.02 | 0.16 | 0.40 |
|  | Temporal | 16.09 (4.21, 0.67) | 17.10 (5.37, 0.80) | 1.01 | 0.480 | 0.21 |
|  | Occipital | 7.87 (2.66, 0.43) | 11.30 (4.46, 0.67) | 3.43 | 0.017 | 0.92 |
| **Theta2** |  |  |  |  |  |  |
|  | Frontal | 13.28 (4.33, 0.69) | 13.65 (4.11, 0.61) | 0.37 | 0.794 | 0.09 |
|  | Central | 13.87 (3.10, 0.50) | 15.26 (3.79, 0.57) | 1.39 | 0.332 | 0.40 |
|  | Parietal | 13.68 (3.89, 0.62) | 14.75 (5.23, 0.78) | 1.07 | 0.455 | 0.23 |
|  | Temporal | 13.22 (3.65, 0.58) | 13.91 (4.39, 0.65) | 0.69 | 0.629 | 0.17 |
|  | Occipital | 10.35 (4.27, 0.68) | 12.65 (4.95, 0.74) | 2.3 | 0.110 | 0.49 |
| **Alpha1** |  |  |  |  |  |  |
|  | Frontal | 24.18 (11.51, 1.84) | 20.65 (12.48, 1.86) | -3.53 | 0.01425 | 0.29 |
|  | Central | 26.78 (12.11, 1.94) | 23.20 (11.80, 1.76) | -3.58 | 0.013 | 0.30 |
|  | Parietal | 31.60 (13.94, 2.23) | 28.12 (12.82, 1.91) | -3.48 | 0.015 | 0.26 |
|  | Temporal | 24.76 (10.84, 1.74) | 22.93 (11.88, 1.77) | -1.83 | 0.203 | 0.16 |
|  | Occipital | 42.29 (17.70, 2.83) | 37.94 (17.52, 2.61) | -4.35 | 0.0025 | 0.25 |
| **Alpha2** |  |  |  |  |  |  |
|  | Frontal | 12.13 (7.43, 1.19) | 7.70 (4.52, 0.67) | -4.43 | 0.0021 | 0.73 |
|  | **Central** | **14.10 (8.20, 1.31)** | **9.25 (4.81, 0.72)** | **-4.85** | **0.00078** | **0.73** |
|  | **Parietal** | **15.63 (9.76, 1.56)** | **10.49 (5.63, 0.84)** | **-5.14** | **0.00035** | **0.66** |
|  | **Temporal** | **14.27 (9.51, 1.52)** | **9.20 (5.10, 0.76)** | **-5.07** | **0.00043** | **0.68** |
|  | **Occipital** | **21.79 (17.73, 2.84)** | **12.08 (7.86, 1.17)** | **-9.71** | **1.76e-11** | **0.73** |
| **Beta1** |  |  |  |  |  |  |
|  | Frontal | 3.34 (1.33, 0.21) | 3.45 (1.59, 0.24) | 0.11 | 0.938 | 0.07 |
|  | Central | 3.44 (1.72, 0.28) | 3.59 (1.78, 0.27) | 0.15 | 0.916 | 0.09 |
|  | Parietal | 3.90 (1.99, 0.32) | 3.95 (2.00, 0.30) | 0.05 | 0.974 | 0.03 |
|  | Temporal | 3.99 (1.93, 0.31) | 3.82 (1.79, 0.27) | -0.17 | 0.905 | 0.09 |
|  | Occipital | 3.03 (2.13, 0.34) | 3.89 (2.72, 0.41) | 0.86 | 0.549 | 0.35 |
| **Beta2** |  |  |  |  |  |  |
|  | Frontal | 2.12 (0.88, 0.14) | 2.63 (3.88, 0.58) | 0.51 | 0.722 | 0.18 |
|  | Central | 2.32 (0.84, 0.13) | 2.37 (1.98, 0.29) | 0.05 | 0.974 | 0.03 |
|  | Parietal | 2.06 (0.84, 0.14) | 2.00 (1.39, 0.21) | -0.06 | 0.968 | 0.05 |
|  | Temporal | 2.23 (0.82, 0.13) | 2.61 (2.77, 0.41) | 0.38 | 0.791 | 0.18 |
|  | Occipital | 1.45 (0.65, 0.10) | 1.73 (1.23, 0.18) | 0.28 | 0.848 | 0.28 |
| **Beta3** |  |  |  |  |  |  |
|  | Frontal | 0.64 (0.39, 0.06) | 0.95 (1.00, 0.15) | 0.31 | 0.829 | 0.40 |
|  | Central | 0.65 (0.33, 0.05) | 0.81 (0.57, 0.09) | 0.16 | 0.912 | 0.34 |
|  | Parietal | 0.51 (0.23, 0.04) | 0.76 (0.73, 0.11) | 0.25 | 0.86 | 0.45 |
|  | Temporal | 0.66 (0.30, 0.05) | 1.17 (1.31, 0.19) | 0.51 | 0.722 | 0.52 |
|  | Occipital | 0.30 (0.16, 0.03) | 0.63 (0.63, 0.09) | 0.33 | 0.818 | 0.70 |
| **Gamma** |  |  |  |  |  |  |
|  | Frontal | 0.17 (0.14, 0.02) | 0.15 (0.12, 0.02) | -0.02 | 0.988 | 0.15 |
|  | Central | 0.18 (0.12, 0.02) | 0.14 (0.11, 0.02) | -0.04 | 0.978 | 0.35 |
|  | Parietal | 0.15 (0.09, 0.01) | 0.13 (0.11, 0.02) | -0.02 | 0.991 | 0.20 |
|  | Temporal | 0.19 (0.13, 0.02) | 0.19 (0.21, 0.03) | 0 | 0.999 | 0.00 |
|  | Occipital | 0.08 (0.04, 0.01) | 0.12 (0.10, 0.01) | 0.04 | 0.982 | 0.51 |

**Figure 1.**
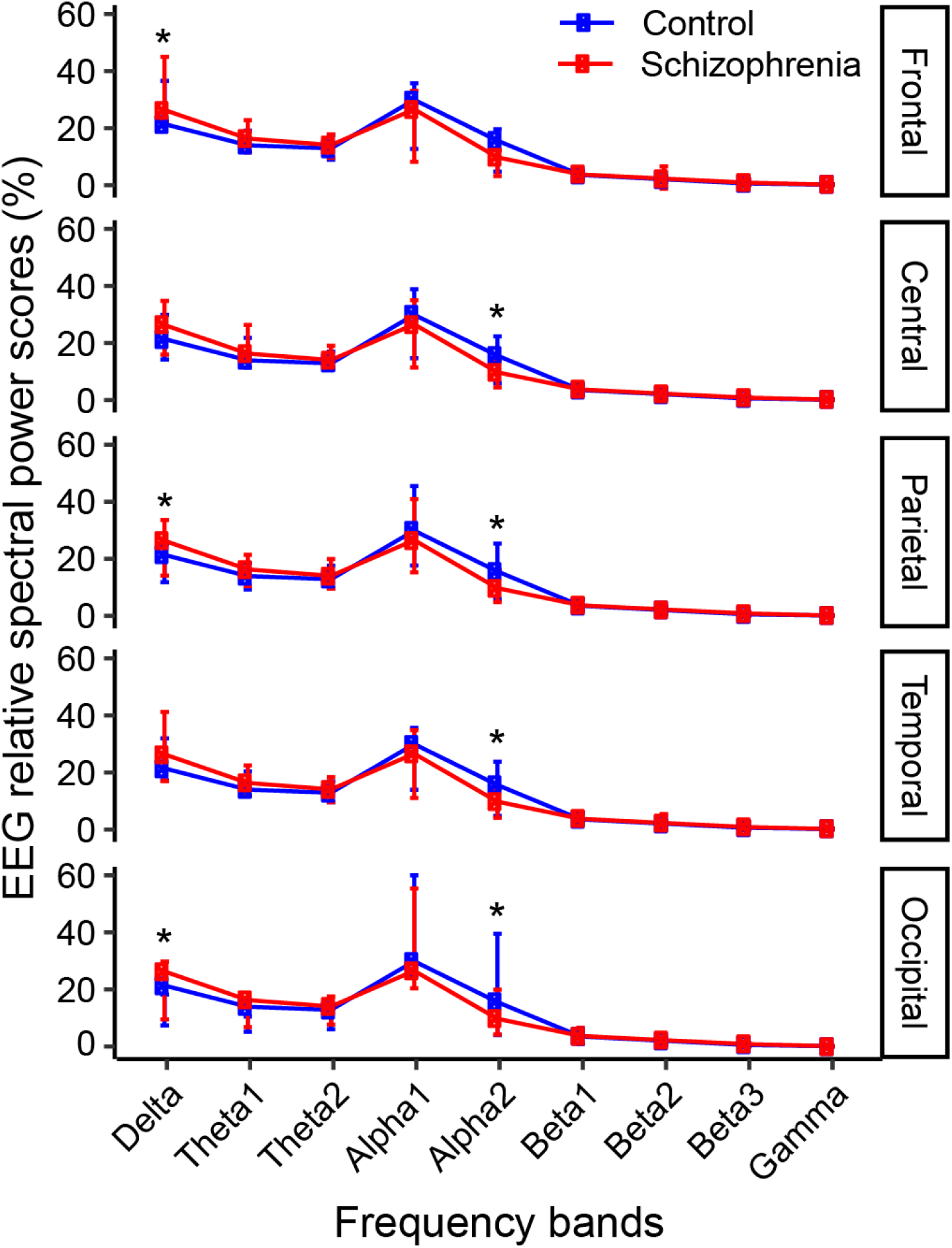
EEG relative spectral power scores (%) for Control (blue) and Schizophrenia (red) groups across nine frequency bands in five cortical regions. Error bars represent ±1 SD. Asterisks denote significance from post-hoc Bonferroni FDR-corrected pairwise comparisons (*p<0.001).

In the delta band, RP was significantly higher in the schizophrenia (SCZ) group than in the control (CON) group. The largest increase was observed in the occipital region (CON: 12.82 ± 5.43% vs. SCZ: 19.65 ± 10.14%; difference = 6.83%; FDR = 2.17 x 10^-6^; |Cohen’s *d*| = 0.82). Significant increases were also observed in the parietal (difference = 5.28%; FDR = 0.0002; |*d*| = 0.62) and the frontal (difference = 4.84%; FDR = 0.00078; |*d*| = 0.49) regions. No significant differences were detected in the central and temporal regions after FDR correction.

In the alpha2 band, RP was significantly lower in the SCZ group than in the CON group. The largest reduction was observed in the occipital region (difference = 9.71%; FDR = 1.76 x 10^-11^; |Cohen’s *d*| = 0.73). Significant reductions were also observed in the parietal (difference = 5.14%; FDR = 0.00035; |*d*| = 0.66), temporal (difference = 5.07%; FDR = 0.00043; |*d*| = 0.68), and central (difference = 4.85%; FDR = 0.00078; |*d*| = 0.73) regions. The frontal region did not meet the FDR-corrected significance threshold.

Across both frequency bands, the occipital cortex exhibited the largest between-group differences, showing the greatest increase in delta RP and the greatest reduction in alpha2 RP (**Table 3**). These findings identified the occipital cortex as the principal region of altered resting-state spectral organization in adolescent schizophrenia.

### 3.4 Topographic Distribution of Relative Power

Topographic maps illustrating the scalp distribution of mean RP for each frequency band are presented in **Figure 2** (left panel: CON; right panel: SCZ). Across most frequency bands, the overall spatial organization of spectral activity was similar between the two groups; however, prominent differences were observed in the delta and alpha2 bands (**Table 3**).

**Figure 2.**
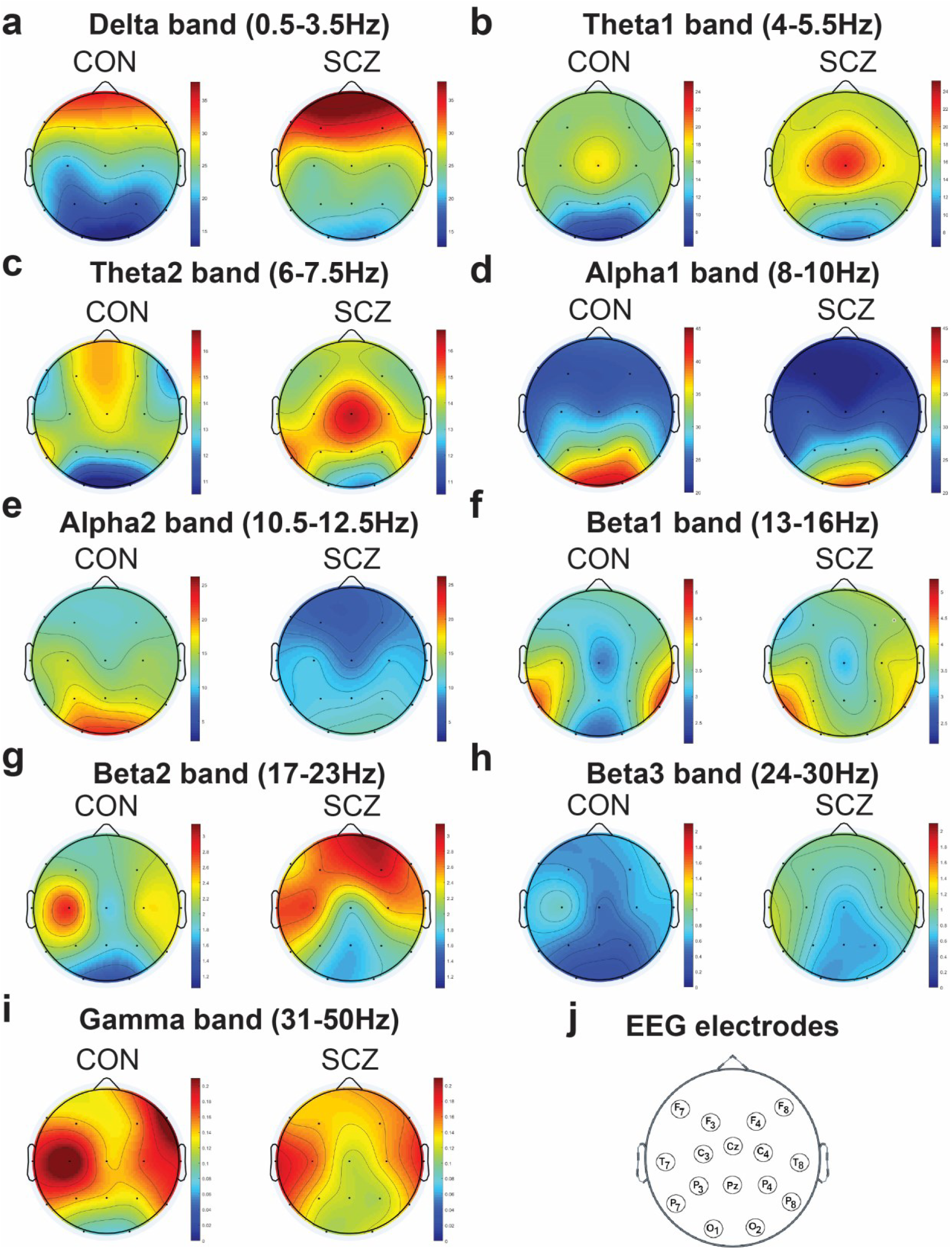
Scalp topographic maps of EEG relative power (%) for each frequency band (a-i). Left: healthy controls (CON, n=39); Right: schizophrenia group (SCZ, n=45). Colour scale represents RP (%). Electrode positions follow the MCN 10-5 system (j).

In the delta band (**Figure 2a**), both groups exhibited a frontal-to-posterior gradient of delta RP. However, the schizophrenia (SCZ) group showed a generalized increase in delta RP across the, with the most pronounced enhancement observed over the posterior cortex, particularly the occipital region. This visual pattern was consistent with the post-hoc analysis, which demonstrated significantly higher delta RP in the frontal, parietal, and occipital regions, with the largest between-group difference observed in the occipital cortex (**Table 3**).

In the alpha bands (**Figures 2d-e**), both groups exhibited the expected occipital predominance of alpha activity. However, the occipito-parietal alpha2 maximum was markedly attenuated in the SCZ group, consistent with the significant reductions in alpha2 RP observed in the occipital, parietal, temporal, and central regions. The largest reduction occurred in the occipital cortex (**Table 3**). In contrast, the alpha1 band showed a similar but less pronounced reduction in posterior alpha activity that did not remain significant after FDR correction (**Table 3**).

The theta (**Figures 2b-c**), beta (**Figures 2f-h**), and gamma (**Figure 2i**) topographies exhibited broadly similar spatial distributions between the SCZ and CON groups, consistent with the absence of robust region-specific differences after FDR correction (**Table 3**).

## 4. Discussion

The present study characterized resting-state EEG relative power (RP) across nine frequency bands and five cortical regions in male adolescents with schizophrenia. We found frequency- and region-specific alterations, including increased delta RP in the frontal, parietal, and occipital cortices and reduced alpha2 RP in the central, parietal, temporal, and occipital cortices, with the largest differences in the occipital region (**Figure 1, Tables 1-3**). Topographic maps visually corroborated these findings by demonstrating widespread posterior delta enhancement together with attenuation of occipito-parietal alpha2 activity (**Figure 2**). Collectively, these findings indicate that resting-state oscillatory activity in adolescent schizophrenia is characterized by selective redistribution of spectral power rather than uniform changes across all frequency bands or cortical regions.

Earlier resting-state quantitative EEG studies primarily reported reduced occipital alpha activity and increased slow-wave predominance using conventional four-band spectral analysis in adults with schizophrenia or schizoaffective disorder^6,7^. In contrast, the present study employed a higher-resolution nine-band spectral decomposition and demonstrated that these abnormalities were predominantly driven by increased delta RP and reduced alpha2 RP, with the most pronounced changes occurring in the occipital cortex of male adolescents with schizophrenia. This approach enabled the identification of alpha2 (10.5-12.5 Hz) as the principal alpha sub-band affected in schizophrenia and demonstrated that the occipital cortex exhibited the most pronounced abnormalities in both delta and alpha2 RP. These findings refine previous observations by providing greater spectral and regional specificity during an early stage of the disorder, before the prolonged effects of chronic illness progression are expected to predominate.

The increase in delta (0.5-3.5 Hz) RP in adolescents with schizophrenia, particularly over the occipital cortex where the largest effect size was detected (FDR: 2.17 x 10^-6^, |Cohen’s *d*|: 0.82; **Table 3**), represented one of the principal findings of this study. Previous resting-state EEG studies have consistently reported elevated slow-wave activity in schizophrenia, predominantly involving the frontal, central, parietal and temporal cortices^3,8,9,24,25^. Similarly, conventional spectral analyses have described occipital alpha reduction together with relative slow-wave predominance in adult schizophrenia and schizoaffective disorder^6,7^. The present findings extend these observations by demonstrating that adolescent schizophrenia is characterized by a marked increase in occipital delta RP, identified through higher-resolution spectral decomposition and quantitative region-wise analysis. Importantly, a systematic review and meta-analysis demonstrated that these slow-wave and alpha abnormalities are robust, largely medication-independent electrophysiological features of schizophrenia, although the cortical topography of slow-wave abnormalities varies across studies^3^. Accordingly, the pronounced occipital delta predominance observed in the present adolescent cohort is consistent with the reported posterior cortical distribution of slow-wave abnormalities in a subset of schizophrenia studies. Collectively, these observations highlight a conserved electrophysiological abnormality with variable regional cortical expression in schizophrenia, as demonstrated in the present adolescent cohort.

Resting-state delta oscillations are generated through coordinated corticothalamic interactions and normally reflect synchronized activity within large-scale cortical networks^26^. Although scalp EEG primarily measures cortical electrical activity, the observed redistribution of spectral power toward slower frequencies may be consistent with disrupted thalamocortical communication proposed in schizophrenia ^4,27^. The pronounced occipital involvement observed in the present study may therefore reflect altered functional interactions between the visual cortex and thalamic relay nuclei, particularly the lateral geniculate nucleus, which forms the principal thalamocortical pathway for visual information processing ^28^. Functional neuroimaging and electrophysiological studies further support widespread abnormalities in thalamocortical connectivity involving visual and frontoparietal networks in schizophrenia^9,24,25^.

Beyond this thalamocortical framing, the schizophrenia-associated slow-wave abnormalities have been linked mechanistically to (a) catechol-O-methyltransferase (COMT) mediated alteration of dopamine metabolism dependent modulation of low-frequency oscillations^29^, (b) impaired occipital-to-frontal bottom-up information transfer resulting from fronto-occipital hierarchical dyscommunication^30^, and (c) N-methyl-D-aspartate receptor (NMDAR) hypofunction-mediated glutamatergic-GABAergic excitation-inhibition imbalance that disrupts synaptic plasticity and large-scale cortical network synchronization^31^. Accordingly, the pronounced occipital delta predominance observed in the present adolescent schizophrenia cohort may reflect the convergent influence of these complementary network- and circuit-level pathophysiological mechanisms.

Because RP represents the proportional distribution of oscillatory activity rather than absolute signal amplitude, the elevated occipital delta RP observed in the present study (**Table 3**; **Figure 1**) most likely reflected a redistribution of resting-state spectral activity (**Figure 2**) toward slower frequencies, rather than an isolated increase in absolute delta power^13^. This interpretation is further supported by the concurrent reduction in occipital alpha2 RP, suggesting a shift in spectral balance (**Figure 2**) rather than independent alterations within individual frequency bands. Together, these findings indicate that posterior cortical oscillatory organization is substantially altered in adolescent schizophrenia and suggest that visual cortical dysfunction may represent an underrecognized neurophysiological feature during the early stages of the disorder. This interpretation is consistent with morphometric evidence demonstrating altered neuronal organization in both the prefrontal cortex (area 9) and primary visual cortex (area 17) in schizophrenia^32^.

The reduction in alpha2 (10.5-12.5 Hz) RP represented another major finding of this study. Significant reductions were observed in the central, parietal, temporal, and occipital cortices, with the largest effect occurring in the occipital cortex (FDR: 1.76×10^-11^, |Cohen’s *d*|: 0.73; **Table 3**). Previous resting-state EEG studies have consistently reported reduced alpha activity in schizophrenia, however, the regional distribution has varied across studies. Conventional four-band analyses described prominent occipital alpha reductions^6,7^, whereas subsequent investigations reported abnormalities involving the frontal, central, parietal, and temporal cortices^8,9^. Despite these observations, most studies analyzed alpha as a broad frequency band, limiting the identification of frequency-specific alterations within the alpha spectrum. By separating alpha oscillations into alpha1 and alpha2 sub-bands, the present study demonstrated that the most robust abnormality was confined to the alpha2 range, indicating that broad-band alpha reductions reported previously may predominantly reflect impairment of higher alpha frequencies.

Alpha oscillations play a central role in sensory gating, cortical inhibition and large-scale neuronal synchronization. Their generation depends on coordinated interactions between thalamocortical circuits and inhibitory cortical interneurons, particularly parvalbumin-positive GABAergic interneurons, which synchronize pyramidal cell firing and regulate rhythmic cortical activity^5^. Dysfunction of these interneurons has been widely implicated in schizophrenia and is thought to contribute to impaired cortical synchronization and reduced alpha oscillatory activity^9,25^. Consequently, the widespread reduction in alpha2 RP observed in the present study (**Figure 1, Table 3**) may reflect impaired inhibitory control within cortical networks, resulting in deficient sensory filtering and abnormal processing of external information.

Reduced occipital alpha power in first-episode psychosis reflects impaired thalamocortical synchronization involving the pulvinar and lateral geniculate nucleus (LGN)^33^, whereas similar abnormalities in clinical high-risk individuals indicate disrupted pulvinar-visual cortical modulation as an early electrophysiological feature of the psychosis spectrum^34^. Schizophrenia is further characterized by downregulation of glutamate decarboxylase 67 (GAD67), reduced GABA receptor function, and impaired GABAergic neurotransmission, molecular abnormalities that compromise inhibitory cortical synchronization and contribute to alpha oscillatory deficits^31^. Consistently, increased occipital alpha functional connectivity despite reduced alpha power in first-episode schizophrenia and ultra-high-risk individuals indicates abnormal reorganization of occipital alpha networks rather than simple attenuation of oscillatory activity^35^. Collectively, these findings indicate that occipital alpha abnormalities emerge early in the psychosis spectrum and reflect impaired posterior thalamocortical network organization. Furthermore, the increased prevalence of visual hallucinations in childhood-onset schizophrenia provides convergent clinical evidence for early involvement of posterior visual cortical systems^36^. Consistent with these observations, the present study extends previous evidence by localizing alpha2 dysfunction predominantly to the occipital cortex with higher spectral and regional resolution, supporting posterior cortical network dysfunction as an early electrophysiological feature of adolescent schizophrenia.

The marked reduction in occipital alpha2 RP study (**Figure 1, Table 3**) further suggests impaired functional organization of the visual cortex during the resting state. Posterior alpha oscillations normally suppress irrelevant visual information and maintain efficient sensory processing, whereas attenuation of these oscillations has been associated with impaired visual perception and abnormal cortical excitability in schizophrenia^25,28^. The concurrent increase in occipital delta RP and reduction in alpha2 RP observed in the present study (**Figure 1, Table 3**) therefore indicate a redistribution of resting-state spectral activity toward slower oscillations within posterior cortical networks. Although scalp EEG (**Figure 2**) does not directly assess thalamic function, this pattern is compatible with previous models proposing disrupted thalamocortical interactions in schizophrenia^4,27^.

The dissociation between alpha1 and alpha2 observed in this dataset is also noteworthy. Although alpha1 exhibited a significant omnibus effect of Diagnosis (**Table 2**), post-hoc analyses did not identify the same pattern of region-specific abnormalities observed for alpha2 after FDR correction. Previous EEG studies rarely distinguished between alpha sub-bands, potentially obscuring frequency-specific alteration^6,7^. Functional evidence suggests that alpha1 and alpha2 reflect partially distinct physiological processes, with alpha2 being more closely associated with attentional control, cognitive processing, and corticocortical communication^37^. The selective reduction of alpha2 RP observed in the present study therefore suggests that higher alpha oscillations may represent a more sensitive electrophysiological feature of adolescent schizophrenia than conventional broad-band alpha measurements.

The scalp topographic maps provided qualitative support for the statistical findings by illustrating the spatial distribution of resting-state oscillatory activity across cortical regions (**Figure 2**). The most prominent differences between groups were observed in the delta and alpha2 frequency bands, with adolescents with schizophrenia exhibiting widespread posterior delta enhancement and attenuation of occipito-parietal alpha2 activity. These visual patterns were consistent with the region-wise statistical analyses and further emphasized the occipital cortex as the principal site of spectral alteration. Because scalp EEG primarily reflects cortical neuronal activity, these findings should be interpreted as evidence of altered cortical oscillatory organization. Nevertheless, the observed cortical abnormalities are compatible with current models proposing disrupted corticothalamic communication and impaired sensory gating in schizophrenia^4,27^.

The present study extends previous resting-state EEG investigations in several important respects. Earlier studies primarily examined adult schizophrenia or schizoaffective disorder using conventional four-band spectral analysis and electrode-wise comparisons^6,7^. In contrast, the present study focused on adolescent schizophrenia and combined a nine-band spectral framework with region-wise quantitative analysis, modern EEG preprocessing, and repeated-measures statistical modelling. This approach demonstrated that alpha2 was the principal alpha sub-band affected and identified the occipital cortex as the region exhibiting the greatest increase in delta RP and the greatest reduction in alpha2 RP. These findings provide greater spectral, regional, and developmental specificity and may improve the neurophysiological characterization of early-onset schizophrenia. Recent multidimensional resting-state EEG analyses indicate that no single EEG feature is fully representative of schizophrenia, as different EEG abnormalities capture distinct aspects of the disorder^38^. Within this framework, the selective occipital delta enhancement and alpha2 attenuation observed in the present study provide greater spectral and regional specificity for characterizing adolescent schizophrenia.

Several limitations of this study should be acknowledged. The dataset was recorded with only 16 EEG channels, limiting spatial resolution compared to high-density arrays. The use of RP rather than AP prevents assessment of whether absolute power increases, decreases, or both underlie the observed spectral redistribution. Finally, the dataset originates from a single Russian research center^10,11^, and cross-cultural and cross-site validation is warranted. Finally, resting-state scalp EEG primarily reflects cortical activity and therefore cannot directly characterize subcortical generators or evaluate the thalamocortical mechanisms proposed by previous neurophysiological models.

## 5. Conclusion

High-resolution analysis of resting-state EEG relative power revealed frequency- and region-specific alterations in male adolescent schizophrenia, characterized by increased delta RP and reduced alpha2 RP, with the most pronounced abnormalities occurring in the occipital cortex. Three principal findings highlight the novelty of the study.

i. **Region-specific increase in delta RP with occipital predominance:** Delta RP was significantly higher in adolescents with schizophrenia in the frontal, parietal and predominantly occipital cortex, but not in the central or temporal regions, revealing a posterior-dominant gradient (occipital > parietal > frontal) that resolves previously inconsistent regional reports of posterior-restricted versus widespread slow-wave abnormalities. As RP reflects proportional spectral distribution rather than absolute amplitude, these findings indicate posterior redistribution of oscillatory power toward slower frequencies rather than increased absolute slow-wave activity.
ii. **Selective alpha2 impairment dissociable from alpha1**: Alpha2 RP was significantly reduced in the central, parietal, temporal, and occipital cortex, representing the largest effect observed, whereas alpha1 showed no region-specific differences after FDR correction. Unlike conventional broad-band alpha analyses, the nine-band subsets decomposition localized the abnormality specifically to alpha2, identifying it as a more sensitive electrophysiological marker than broad alpha.
iii. **Occipital cortex as the principal locus of spectral redistribution**: The occipital cortex exhibited the largest effects in both significant frequency bands (delta and alpha2), with topographic maps confirming posterior delta enhancement and occipito-parietal alpha2 attenuation. The band-specific differences, despite the absence of overall regional effects, demonstrate that schizophrenia-related EEG abnormalities converge on the occipital cortex in a frequency-specific manner rather than reflecting diffuse cortical dysfunction.

These findings extend previous resting-state EEG studies by providing greater spectral and regional specificity in adolescent schizophrenia and suggest that EEG-derived relative power may serve as a promising neurophysiological feature for characterizing early-onset schizophrenia.

## Acknowledgements

The authors thank the CPEPA-UGC Centre for “Electrophysiological and Neuro-imaging Studies including Mathematical Modeling and Department of Physiology, University of Calcutta, for institutional support and research facilities. They also thank the clinicians and researchers at the National Center of Mental Health of the Russian Academy of Medical Sciences and Moscow State University for providing public access to the male adolescent schizophrenia EEG dataset.

## CRediT authorship contribution statement

**Suvojit Hazra:** Conceptualization, Data curation, Formal analysis, Investigation, Methodology, Software, Validation, Visualization, Writing – original draft. **Nilkanta Chakrabarti:** Conceptualization, Project administration, Resources, Supervision, Validation, Writing – review & editing.

## Data availability

All the data are presented in the article.

